# Similarity judgments and cortical visual responses reflect different properties of object and scene categories in naturalistic images

**DOI:** 10.1101/316554

**Authors:** Marcie L. King, Iris I. A. Groen, Adam Steel, Dwight J. Kravitz, Chris I. Baker

**Author notes:** co-first authors (Marcie L. King, Iris I. A. Groen). Corresponding author: CIB.

## Abstract

Numerous factors have been reported to underlie the representation of complex images in high-level human visual cortex, including categories (e.g. faces, objects, scenes), animacy, and real-world size, but the extent to which this organization is reflected in behavioral judgments of real-world stimuli is unclear. Here, we compared representations derived from explicit similarity judgments and ultra-high field (7T) fMRI of human visual cortex for multiple exemplars of a diverse set of naturalistic images from 48 object and scene categories. Behavioral judgements revealed a coarse division between man-made (including humans) and natural (including animals) images, with clear groupings of conceptually-related categories (e.g. transportation, animals), while these conceptual groupings were largely absent in the fMRI representations. Instead, fMRI responses tended to reflect a separation of both human and non-human faces/bodies from all other categories. This pattern yielded a statistically significant, but surprisingly limited correlation between the two representational spaces. Further, comparison of the behavioral and fMRI representational spaces with those derived from the layers of a deep neural network (DNN) showed a strong correspondence with behavior in the top-most layer and with fMRI in the mid-level layers. These results suggest that there is no simple mapping between responses in high-level visual cortex and behavior – each domain reflects different visual properties of the images and responses in high-level visual cortex may correspond to intermediate stages of processing between basic visual features and the conceptual categories that dominate the behavioral response.

**Significance Statement:** It is commonly assumed there is a correspondence between behavioral judgments of complex visual stimuli and the response of high-level visual cortex. We directly compared these representations across a diverse set of naturalistic object and scene categories and found a surprisingly and strikingly different representational structure. Further, both types of representation showed good correspondence with a deep neural network, but each correlated most strongly with different layers. These results show that behavioral judgments reflect more conceptual properties and visual cortical fMRI responses capture more general visual features. Collectively, our findings highlight that great care must be taken in mapping the response of visual cortex onto behavior, which clearly reflect different information.

## Introduction

The ventral visual pathway, extending from primary visual cortex (V1) through the inferior temporal lobe, is thought to be critical for object, face and scene recognition (Kravitz et al., 2013). While posterior regions in this pathway respond strongly to the presentation of low-level visual features, more anterior regions are thought to encode high-level categorical aspects of the visual input. For example, functional magnetic resonance imaging (fMRI) studies have identified category-selective regions in ventral temporal cortex (vTC) and lateral occipitotemporal cortex (lOTC) that show preferential responses for images of one category compared to another (e.g. face-selective fusiform face area or FFA, scene-selective parahippocampal place area or PPA, and object-selective lateral occipital complex or LOC; Kanwisher and Dilks, 2013). However, many other factors have been reported to contribute to responses in high-level visual cortex, including, but not limited to, eccentricity (Hasson et al., 2003), elevation (Silson et al., 2015), real-world size (Konkle and Oliva, 2012), typicality (Iordan et al., 2016), category level (i.e. superordinate, basic, subordinate – Iordan et al., 2015), and animacy (Kriegeskorte et al., 2008; Connolly et al., 2012; Naselaris et al., 2012; Sha et al., 2015; Proklova et al., 2016). The goal of the current study was determine the correspondence between the response of high-level visual cortex and our mental representations of category by comparing the representational space reflected in fMRI responses with behavioral similarity judgements for naturalistic images across a broad range of object and scene categories.

Determining how responses in high-level visual cortex relate to behavior is critical for elucidating the functional significance of these regions. For tasks such as identification and categorization, relevant information has been reported in the responses of lOTC and vTC (Kravitz et al., 2013; Grill-Spector and Weiner, 2014) and it is commonly assumed there is a direct mapping between responses in high-level visual cortex and behavioral judgments. But this assumption belies the diverse behavioral goals these regions likely support (Malcolm et al., 2016; Peelen and Downing, 2017). While the fMRI responses in both human and non-human primate vTC appear to reflect major distinctions between animate/inanimate and face/body, behavioral similarity judgements reveal additional fine-grained representational structure, patricularly for inanimate objects (Kriegeskorte et al., 2008; Mur et al., 2013). However, these studies contained a limited sampling of different categories that emphasized some categories (e.g. faces, food/fruit) over others (e.g. chairs, appliances) and may have only captured part of the representational structure. While other fMRI studies have included a broader sampling of different categories (Huth et al., 2012; Naselaris et al., 2012), behavioral judgments were not collected beyond labels for discrete elements of the images that may not characterize the broader conceptual representation. Here, we combined a varied sampling of different categories with both ultra-high field (7T) fMRI and detailed behavioral similarity measurements to determine what aspects of representation are shared between behavior and the response of high-level visual cortex.

We presented multiple images from 48 categories ranging across both object (e.g. bags, dolls) and scene (e.g. kitchens, mountains) categories. In contrast to some prior studies that presented segmented objects with limited, arbitrary or no context (Kriegeskorte et al., 2008; Konkle and Oliva, 2012; Yamins et al., 2014) our study used objects in typical contexts. We found highly reproducible but distinct structure in both behavior and fMRI with little evidence for the previously reported animacy division. Instead, behavioral judgments reflected a manmade/natural division, while cortical regions largely showed a separation of images containing human and non-human faces and bodies from everything else. Computational features extracted from a deep neural network (DNN) trained on object recognition correlated with representational structure in both behavior and fMRI, but the strongest match with behavior was with the highest DNN layer, while fMRI correlated best with a mid-level DNN layer. Collectively, these results suggest that while both behavior and the response of high-level visual cortex reflect combinations of visual features, those features differ between domains, with no direct mapping between them.

## Materials and Methods

### Stimuli

We retrieved high-resolution (1024×768 pixels) color photographs from Google Images to construct two sets of stimuli, each comprised of 144 individual color images of complex scenes. We included two separate sets to be able to test generalization of our findings across images. Each set of images (hereby referred to as Image Set 1 and Image Set 2) contained 48 concrete categories, with 3 exemplar images per category (Figure 1). The 48 categories were chosen to reflect a diverse range of naturalistic object and scene categories. All of the images in Image Set 1 and Image Set 2 depicted people, places, and things in natural context and from familiar viewpoints. The images portrayed scenes that one might expect to see on a typical day, and were chosen for their neutral nature (i.e. to be unlikely to elicit any strong emotional response).

**Figure 1:**
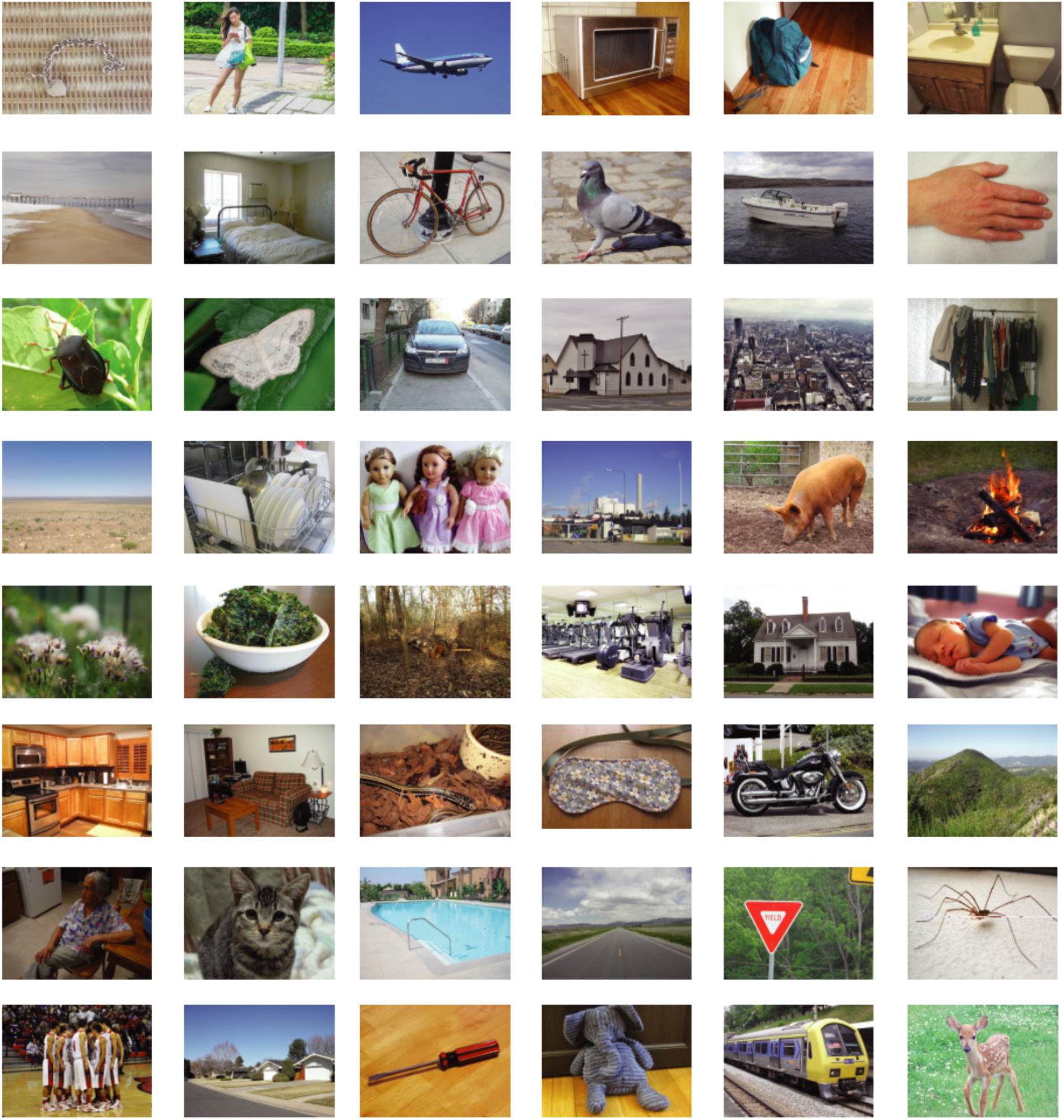
Naturalistic image categories. One exemplar from each of the 48 image categories, presented in alphabetical order: accessories, adults, airplanes, appliances, bags, bathrooms, beaches, beds, bikes, birds, boats, body parts, bugs, butterflies, cars, churches, cityscapes, clothes, deserts, dishes, dolls, factories, farm animals, fire, flowers, food, forests, gyms, houses, kids, kitchens, living rooms, lizards/snakes, masks, motorcycles, mountains, older adults, pets, pools, roads, signs, spiders, sports, suburbs, tools, toys, trains, wild animals.

### Participants and testing

20 healthy human volunteers (9 male, mean age = 27.7 years) participated in the behavioral similarity judgment experiment. 10 participants viewed Image Set 1 (4 male, mean age = 29.3) and 10 participants viewed Image Set 2 (5 male, mean age = 26.1). 10 of these participants also participated in the corresponding fMRI experiment prior to participating in the behavioral portion of this study. 5 of these participants viewed stimuli from Image Set 1 (3 male, mean age = 26.6 years) and 5 participants viewed stimuli from Image Set 2 (2 male, mean age = 26.2 years). Each participant saw the same stimulus set in both the behavioral and fMRI experiment. All fMRI participants completed the fMRI scan session before rating the behavioral similarities of the images. This study was conducted in accordance with The National Institutes of Health Institutional Review Board, and all participants gave written informed consent as part of the study protocol (93 M-0170, NCT00001360) prior to participation in the study.

### Behavioral paradigm

We adopted a multi-arrangement paradigm previously used by Kriegeskorte, Mur and colleagues (Kriegeskorte and Mur, 2012; Mur et al., 2013). Participants were seated at a distance of approximately 50 cm in front of a computer monitor (Dell U3014, 30 inches, 2560 × 1600 pixels) and completed the object arrangement task on 144 images comprising either Image Set 1 or Image Set 2. At the onset of the task, all 144 images were presented simultaneously in random order around the perimeter of a circle presented on the computer monitor, forming an “arena” in which similarity judgments were made. Participants were instructed to “please arrange these images according to their similarity, whatever that means to you. Images that are more similar should go closer together and images that are less similar should go farther apart.” These instructions were purposefully general so as not to bias the arrangements of the images in any particular way, allowing us to investigate what dimensions participants spontaneously use when judging the similarity between images. Participants dragged the individual images into the arena using the mouse and physically arranged them according to their perceived similarity. Given the large number of images (and thus the small size each could be presented at), when a participant clicked on a particular image in the arena, an enlarged version of the image (150 × 200 pixels) was displayed in the top right of the computer screen.

Given the large number of images in the stimulus sets, participants completed only one arrangement of the images, in contrast to the original implementation of this method that used additional trials with selective subsets of stimuli (Kriegeskorte et al., 2012). However, participants were able to re-arrange images within the circular area on the screen after their initial placement as many times as they wanted within a 1-hour time limit, and they were encouraged to verify that they were satisfied with the final arrangement. In addition, in our experience this task exhibits very high correlations between results of the first and the last trial (unpublished data). One of the benefits of this arrangement method is that we were able to collect a large number of simultaneous pairwise similarity judgments in a reasonably short amount of time. Perceived object-similarity is traditionally measured using pairwise similarity judgments, however it would take many hours and testing sessions to acquire judgments on our 10,296 possible pair combinations of images. Therefore, in the current method we used the spatial arrangement of the images as a measure of their perceived similarity. Specifically, the Euclidean distance between an image and every other image was used as the measurement of perceived dissimilarity between the images (i.e. dissimilarity estimate). Representational dissimilarity matrices (RDMs) were constructed for each participant, using the ranked dissimilarity estimates for each image pair. Note that the distance matrix discards the absolute position of stimuli and only retains their relative location, which should minimize bias related to the initial placement of the stimuli.

### fMRI paradigm

Participants were scanned while viewing the stimuli on a back-projected screen through a rear-view mirror that was mounted on the head coil. Stimuli were presented at a resolution of 1024 × 768 pixels and subtended 20 × 15 degrees of visual angle. Individual scenes were presented in an event-related design for a duration of 500 ms, separated by a 5 s interval. Throughout the experimental run, a small fixation cross (<0.5 degrees) was presented in the center of the screen. Participants viewed all 144 images in either Image Set 1 or Image Set 2 while performing an unrelated fixation cross task. Simultaneous with the onset of each stimulus, either the vertical or horizontal arm of the fixation cross became slightly elongated. Participants were asked to indicate, via button response, whether the horizontal or vertical line of the fixation cross was longer. Both arms changed equally often within a given run, and arm changes were randomly assigned to individual stimuli. Participants completed 12 runs of the event-related experiment, with each run being composed of 156 TRs. Within each run, 48 images were presented such that after 3 consecutive runs participants had viewed the entire set of 144 images. Thus, participants viewed 4 complete repeats of the 144 images in total.

### Scanning parameters

Participants were scanned on a research-dedicated Siemens 7 Tesla Magnetom scanner in the Clinical Research Center on the National Institutes of Health campus in Bethesda, Maryland. Partial T2^*^-weighted functional image volumes of the frontal, temporal, and occipital cortices were acquired using a 32-channel head coil (47 slices; 1.6 × 1.6 × 1.6 mm isotropic voxels; 10 % interslice gap; TR 2 s; TE 27 ms; flip angle 70°, matrix size 126 × 126; FOV 200 mm). In all scans, oblique slices were oriented approximately parallel to the ventral portion of the prefrontal cortex. In addition, standard MPRAGE (magnetization-prepared rapid-acquisition gradient echo) and corresponding GE-PD (gradient echo–proton density) images were acquired, and the MPRAGE images were then normalized by the GE-PD images for use as a high-resolution anatomical image for the following fMRI data analysis (Van de Moortele et al., 2009).

### Functional localizers

During each scan session, an independent functional localizer scan was also collected in each participant to identify scene and face selective regions in ventral temporal and lateral occipitotemporal cortex. The localizer used an on-off design, alternating between 16 s blocks of scene images and blocks of face images presented at 5 × 5° of visual angle. Localizer runs comprised 144 TRs. Participants performed a one-back task, responding to immediate repeats of the same image using a button press.

### fMRI data preprocessing

All imaging data were processed using the Analysis of Functional NeuroImages (AFNI) software package (http://afni.nimh.nih.gov/afni, RRID:SCR_005927). Prior to statistical analyses, the functional scans were slice-time corrected and all images were motion corrected to the first image of the first functional run, after removing the appropriate number of ‘dummy’ volumes (6) to allow for stabilization of the magnetic field. Following motion-correction, data were smoothed with a 2 mm full-width at half-maximum Gaussian kernel.

### Functionally defined ROIs

Scene and face selective regions of interest (ROIs) were created for each participant based on the localizer runs. A response model was built by convolving a standard HRF function with the block structure for each run and was correlated to the activation time course. ROIs were generated by thresholding the statistical parametric maps at a threshold of *p* < 0.0001 (uncorrected). Contiguous clusters of voxels (> 20) exceeding the defined threshold were defined as scene or face selective. The anatomical locations of these clusters were then inspected to ensure that the current ROIs were consistent with those described in previously published work (Kanwisher, 2010). Our functionally defined face-selective regions included the Fusiform Face Area (FFA) and Occipital Face Area (OFA), and our functionally defined scene-selective regions included the Parahippocampal Place Area (PPA) and the Occipital Place Area (OPA). Ventral early visual areas (vEVC) and dorsal early visual (dEVC) areas (V1-V3) were defined using previously acquired retinotopic field maps from independent participants (Silson et al., 2015, 2016a).

### Anatomically defined ROIs

Anatomically defined ROIs were constructed using the Freesurfer image analysis suite, which is documented and freely available for download online (http://surfer.nmr.mgh.harvard.edu/). A ventral temporal cortical (vTC) region was defined using the lower edge of the inferior temporal sulcus as the lateral boundary, extending medially to include the collateral sulcus. Posteriorly, the vTC extended to the edge of the EVC ROIs and anteriorly to the tip of the collateral sulcus This vTC ROI overlapped with both the functionally-defined FFA and PPA and was drawn to be analogous to the human IT ROI used by Kriegeskorte and colleagues (Kriegeskorte et al., 2008). In addition, a lateral occipitotemporal (lOTC) region was defined extending from the junction of the dorsal and ventral EVC ROIs anteriorly to the superior temporal sulcus, superiorly to the intraparietal sulcus and ventrally to the inferior temporal sulcus. This lOTC ROI overlapped with both the functionally-defined OFA and OPA and also included retinotopic regions such as V3A, LO1 and LO2 (Larsson and Heeger, 2006).

### fMRI analysis: event-related data

All 12 functional runs were concatenated and compared to the activation time course for each stimulus condition using Generalized Least Squares (GLSQ) regression in AFNI. In the current paradigm, each image was treated as an independent condition, resulting in 144 separate regressors for each individual stimulus condition, as well as motion parameters and four polynomials to account for slow drifts in the signal. To derive the response magnitude per stimulus, t-tests were performed between the stimulus-specific beta estimates and baseline for each voxel. All subsequent analyses of these data were conducted in Matlab (<u>Mathworks, Natick</u>, RRID:SCR_001622). To derive representational dissimilarity matrices (RDMs), pairwise Pearson’s correlations were computed between conditions using the t-values across all voxels within a given ROI (Kravitz et al., 2010, 2011). The resulting RDM for a given ROI was a 144 × 144 matrix representing the pairwise correlations between patterns of activity elicited by each stimulus condition. RDMs were created for each participant, ranked using a tied ranking procedure, and then averaged together across participants for each ROI.

### Behavior-fMRI comparisons

We calculated full correlations between behavioral judgment RDMs and each of the fMRI derived RDMs (Spearman’s *ρ)*. For all analyses, the behavioral RDMs were based on averages across the maximum number of participants available for that analysis (e.g., all 20 subjects that performed the behavioral experiment for the group-average behavioral judgments; all 10 subjects that performed the behavioral task on Image Set 1 for the group average behavioral RDM of Image Set 1), with the exception of the within-subject behavior-fMRI comparisons (Figure 4) in which only the participants that also performed the fMRI experiment were included. Statistical significance of between-RDM correlations was determined using fixed-effects stimulus-label randomization tests (Nili et al., 2014). For these tests, a null distribution of between-RDM correlations was obtained by permuting stimulus condition labels of one of the subject-averaged RDMs (e.g., behavioral RDM) 10,000 times, after which the p-value of the observed correlation was determined as its two-tailed probability level relative to the null distribution. In addition, 95% confidence intervals and standard deviations were determined using bootstrap resampling, whereby a distribution of correlation values was obtained by sampling stimulus conditions with replacement (n = 10,000 bootstraps). To correct for multiple testing of the behavioral RDM against the multiple fMRI ROIs, the resulting *p*-values were corrected for multiple comparisons across all ROIs using FDR-correction at α = 0.05.

### Hierarchical clustering

To reveal higher-order relations between the image categories, the behavioral and fMRI measurements were subjected to hierarchical clustering. To estimate the number of clusters that best described the data, we performed k-means clustering (‘kmeans’ function implemented in Matlab, 28 iterations) and evaluated the trade-off between number of clusters and explained variance using the elbow method. Using this method, we determined that six clusters optimally described the behavioral data (80% variance explained in each image set). We subsequently performed hierarchical clustering on both the behavioral judgement RDMs and fMRI derived RDMs (‘cluster’ function in Matlab, method: ‘linkage’, number of clusters: 6).

### Searchlight analysis

To test the relationship between behavioral similarity judgments and activity recorded outside specified ROIs, we conducted whole-brain searchlight analysis. The searchlight analysis stepped through every voxel in the brain and extracted the t-values from a sphere of 3 voxel radius around that voxel (total number of voxels per searchlight sphere = 123), which were then used to compute pairwise correlation distances (1-Pearson’s *r*) between each stimulus condition. Analogous to the ROI analyses, the resulting RDMs were correlated (Spearman’s *rho*) with the average behavioral RDM. These correlation coefficients were assigned to the center voxel of each searchlight, resulting in a separate whole-volume correlation map for each participant computed in their native volume space. To allow comparison at the group level, individual participant maps were first aligned to their own high-resolution anatomical T1 and then to surface reconstructions of the grey and white matter boundaries created from these T1s using the Freesurfer (http://surfer.nmr.mgh.harvard.edu/, RRID:SCR_001847) 5.3 autorecon script using SUMA (Surface Mapping with AFNI) software (https://afni.nimh.nih.gov/Suma). Group-level significance was determined by submitting these surface maps to node-wise *t*-tests in conjunction with Threshold Free Cluster Enhancement (Smith and Nichols, 2009) to correct for multiple comparisons, using the CoSMoMVPA toolbox (Oosterhof et al., 2016).

### DNN comparisons

Deep convolutional neural networks (DNNs) are state-of-the-art computer vision models capable of labeling objects in natural images with human-level accuracy (Krizhevsky et al., 2012; Kriegeskorte, 2015), and are therefore considered potentially relevant models of how object recognition may be implemented in the human brain (Kriegeskorte, 2015; Yamins and DiCarlo, 2016; Scholte, 2017; Tripp, 2017). DNNs consist of multiple layers that perfom transformations from pixels in the input image to a class label through a non-linear mapping of local convolutional filters responses (layers 1–5) onto a set of fully-connected layers of classification nodes (layers 6–8) culminating in a vector of output ‘activations’ for labels assigned in the DNN training phase. Inspection of the learned feature selectivity (Zhou et al., 2014; Güçlü and van Gerven, 2015; Bau et al., 2017; Wen et al., 2017) show that earlier layers contain local filters that resemble V1-like receptive fields while higher layers develop selectivity for entire objects or object parts, perhaps resembling category-selective regions in visual cortex. The feature representations learned by these DNNs have indeed been shown to exhibit some correspondence with both behavior and brain activity measurements in humans and non-human primates during object recognition (Khaligh-Razavi and Kriegeskorte, 2014; Yamins et al., 2014; Güçlü and van Gerven, 2015; Cichy et al., 2016) and scene recognition (Greene et al., 2016; Bonner and Epstein, 2017; Martin Cichy et al., 2017; Groen et al., 2018).

We used the MatConvNet toolbox (Vedaldi and Lenc, 2015) to implement a pre-trained version of an 8-layer deep convolutional neural network (VGG-S CNN) (Chatfield et al., 2014) that was trained to perform the 1000-class ImageNet ILSVRC 2012 object classification task. DNN representations for each individual image in both stimulus sets were extracted from layers 1-5 (convolutional layers) and 6-8 (fully-connected layers) of the network. For each layer, we calculated the Pearson correlation coefficient between each pairwise combination of stimuli yielding one 144 × 144 RDM per DNN layer. Analogous to the behavior-fMRI analyses, we then calculated Spearman’s rank correlations between RDMs derived from DNN layers and RDMs derived from the fMRI and behavioral measurements. Statistical significance was again determined using stimulus-randomization (n = 10,000 permutations, two-tailed tests). Differences in correlation between individual layers were determined using bootstrap tests (n = 10,000) whereby the p-value of a difference in correlation between two layers was estimated as the proportion of bootstrap samples further in the tails (two-sided) than 0 (Nili et al., 2014). To correct for multiple testing of several model representations against the same RDM, the resulting p-values were corrected for multiple comparisons across all tests conducted for a given behavioral or fMRI RDM using FDR-correction at α = 0.05.

## Results

The primary aim of this study was to elucidate the representational space of complex naturalistic categories as reflected in human behavior and in neural responses measured with fMRI. We first present analyses examining and comparing the representational structure of each image set estimated from both behavioral similarity judgments and from fMRI responses in visual cortex. We then examine to what extent features derived from a deep neural network (DNN) model can explain the behavioral and fMRI data.

### Comparison of behavioral judgments and fMRI: Representational Dissimilarity Matrices (RDMs)

We first created RDMs based on both the behavioral judgments and fMRI responses, separately for Image Set 1 and Image Set 2. For behavioral judgments, dissimilarities were based on the pixel distances between images in the multi-arrangement similarity task. For fMRI, we focused on the pairwise comparisons of multi-voxel patterns for each stimulus in ventral temporal cortex using a vTC ROI following Kriegeskorte and colleagues (Kriegeskorte et al., 2008; see Methods). The resulting RDMs are organized alphabetically by category (Figure 2).

**Figure 2:**
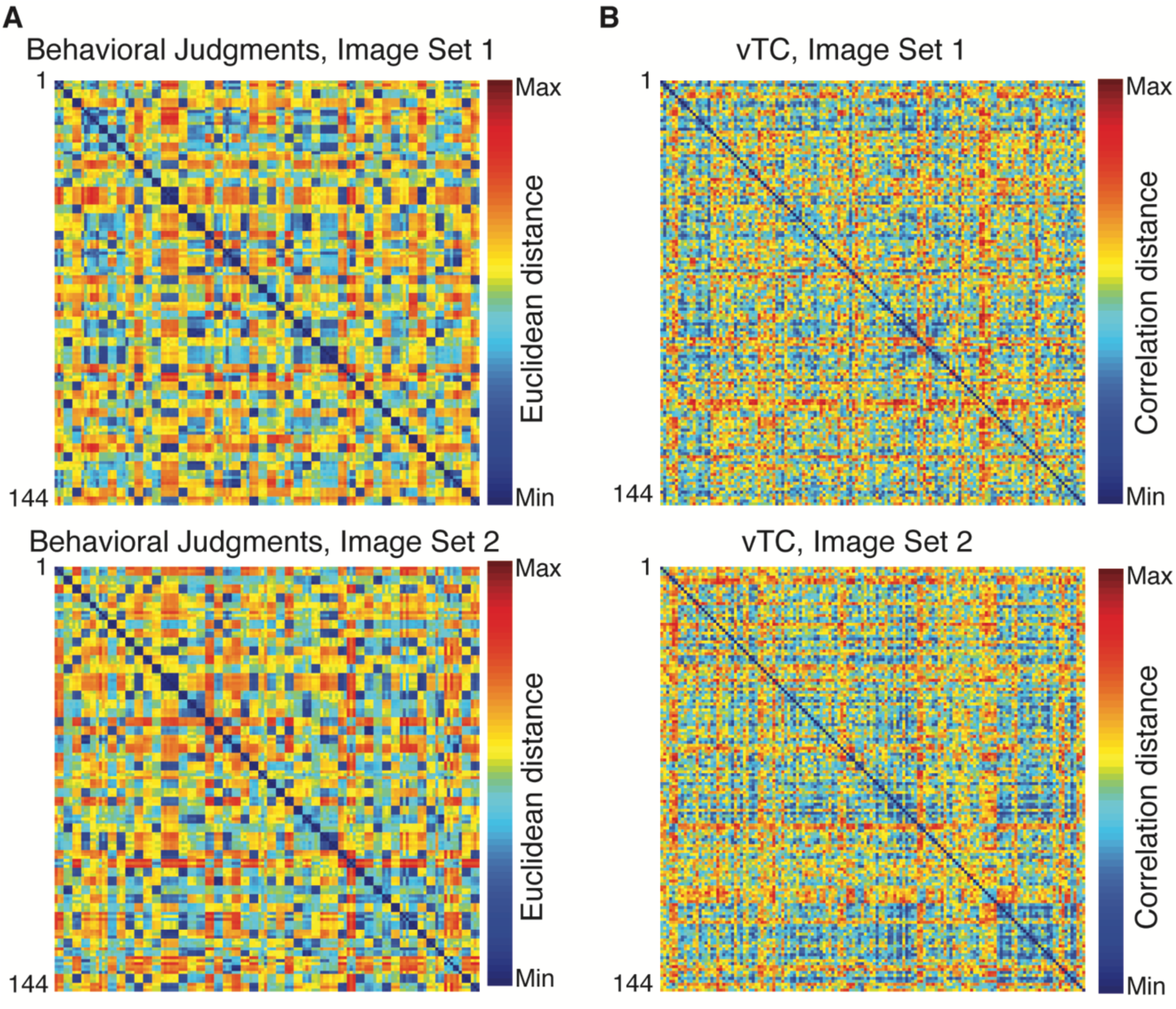
Representational dissimilarity matrices for Image Set 1 and Image Set 2. Matrices show comparisons for all 144 images grouped alphabetically by category (3 images per category, same order as Figure 1). A) Behavioral dissimilarity was measured as the Euclidean distance between pairs of images in the multi-arrangement task. Clustering-by-category is evidenced by the appearance of 3 × 3 exemplar ‘blocks’ exhibiting low dissimilarity along the diagonal. B) fMRI dissimilarity was measured as 1 minus the pairwise correlation between the pattern of response to images in vTC. There is some clustering-by-category present, but it is less evident than for the behavioral judgments.

For behavioral judgments, these RDMs exhibit a clear clustering of exemplars within each category for both image sets (Figure 2A). Participants judged exemplars of the same category as more similar to other exemplars within the same category than to exemplars in different categories (e.g. body parts are more similar to body parts than to mountains). In contrast, there was much less clustering of exemplars for the vTC RDMs, even within category (Figure 2B). The striking difference between behavioral and fMRI RDMs is reflected in weak, albeit significant, correlations between the two measures (Image Set 1, *rho* = 0.06, 95% CI = [0.02, 0.14], *p* = 0.012; Image Set 2, *rho* = 0.07, CI = [0.03, 0.15], *p* = 0.004), suggesting limited similarity in the representation of the images at the image level in behavioral similarity judgements and vTC.

To quantify the extent of category coherence in each image set, we calculated a Category Index as the difference between the average within-category distance and the average between-category distance (Figure 3A). For both behavioral judgments and vTC, this Category Index was greater than zero for both image sets (behavior Image Set 1: one-sample t-tests: *t*(47) = 41.6, CI = [0.40, 0.45], *p* < 0.0001; behavior Image Set 2: *t*(47) = 44.3, CI = [0.42, 0.46], *p* < 0.0001; vTC Image Set 1: *t*(47) = 5.2, CI = [0.05, 0.11], *p* < 0.0001; vTC Image Set 2: *t*(47) = 6.3, CI = [0.05, 0.09], *p* < 0.0001), indicating the presence of significant categorical structure in both domains. However, categorization was much stronger for the behavioral judgments compared to vTC (independent samples t-test: *t*(94) = 29.7, CI = [0.34, 0.39], *p* < 0.001).

**Figure 3:**
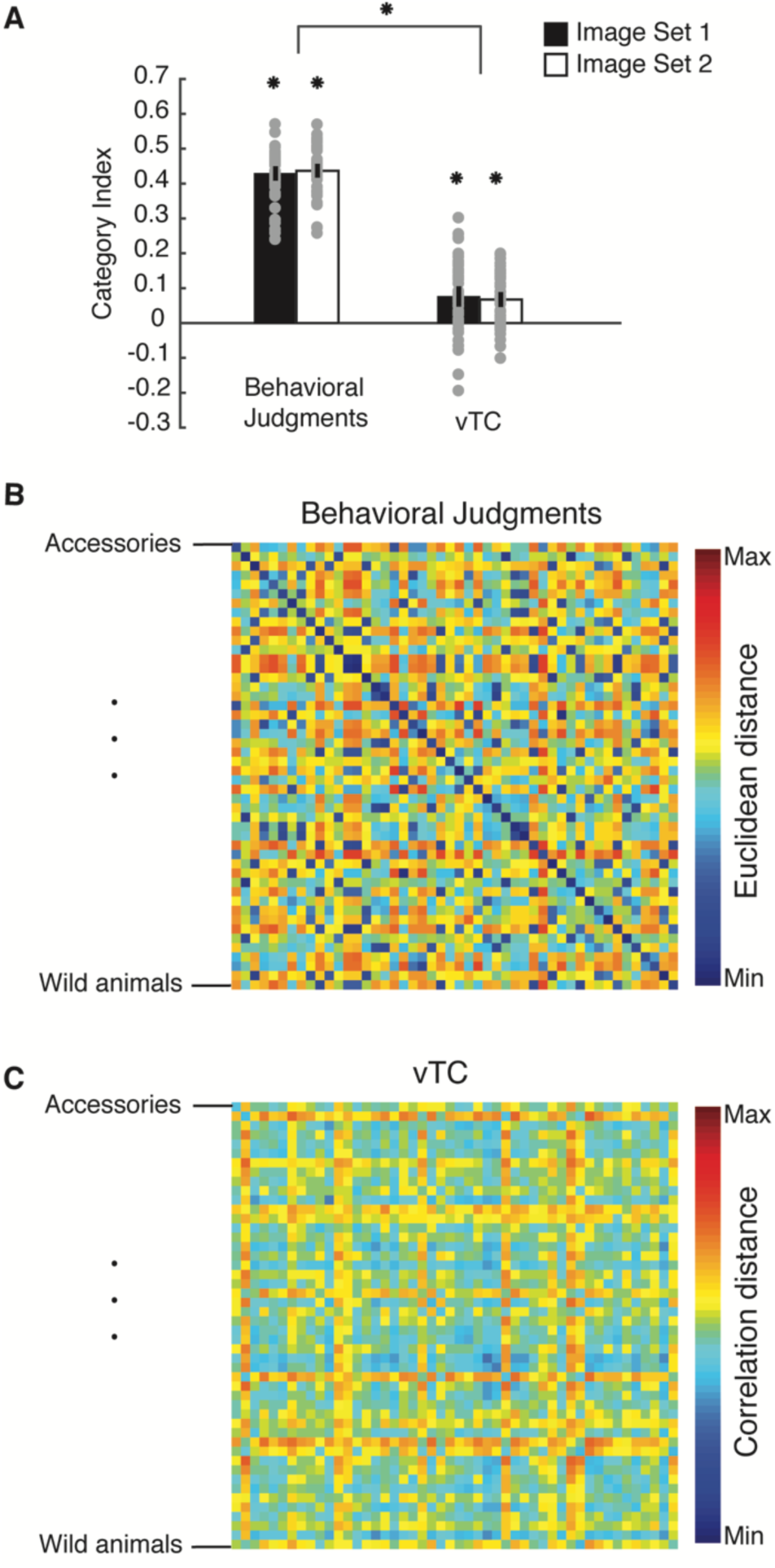
Category representations. a) Category indices for vTC and behavioral similarity judgements calculated as the difference between the average within-category and between-category distances, averaged across categories. Gray dots indicate indices for each category separately. Error bars indicate 95% confidence intervals estimated from a one-sample *t*-test. ^*^ = *p* < 0.001. b), c) RDMs averaged by category for behavioral similarity judgements and fMRI responses in vTC. Categories are ordered alphabetically in the matrices.

Given the presence of significant categorical structure in both domains, and to directly compare Image Set 1 and Image Set 2, which contained different exemplars for each category, we averaged across exemplars (excluding the diagonal), reducing our 144 × 144 exemplar-level RDMs to 48 × 48 category-level RDMs. For both behavioral judgments and vTC there was a strong positive correlation between Image Set 1 and Image Set 2 (behavioral judgments, *rho* = 0.64, CI = [0.55, 0.76], *p* < 0.0001; vTC, *rho* = 0.48, CI = [0.32, 0.67], *p* < 0.0001), indicating that the representational structure in both domains is reproducible across image sets.

Given this reproducibility of representational structure across image sets in both behavior and vTC, we averaged across sets to compare the representational space at a category-level between behavior and vTC (Figures 3B, C). Similar to the exemplar level, there was only a weak, albeit significant, correlation between behavioral judgments and vTC (*rho* = 0.10, CI = [0.02, 0.32], *p* = 0.019). Notably, this correlation was weaker than the relationship between Image Set 1 and Image Set 2 within behavior and vTC separately (Fisher’ r to z transformation: behavior-vTC correlation vs. behavior-behavior Image Set correlation: *z*(48) = 3.1, *p* = 0.002 (two-tailed); behavior-vTC correlation vs. vTC-vTC Image Set correlation: *z*(48) = 2.0, *p* = 0.045 (two-tailed)). Thus, at both the exemplar and category level there was only weak agreement between the representational structure reflected in behavioral judgments and that derived from vTC, despite reliable representational structure across image sets for both behavioral judgments and vTC.

The difference between the representational structure in behavior and vTC may be due to greater variation in the structure across individuals. To address this question, we compared the representational structure from behavior and vTC of the individual participants (Figure 4). This analysis was consistent with the group-level findings: in general, across participants, correlation within an experimental measure (behavior, vTC response) was greater than zero (behavior: range *rho* = [0.05, 0.47]; vTC: range *rho* = [−0.02, 0.41]), suggesting that within a domain the structure of representation was consistent across individuals. However, between experimental measures, correlations were weaker (range *rho* = [−0.06, 0.18]), even for the same participant. Thus, there was not a strong relationship between a single participant’s behavioral RDM and his or her own vTC RDM.

**Figure 4:**
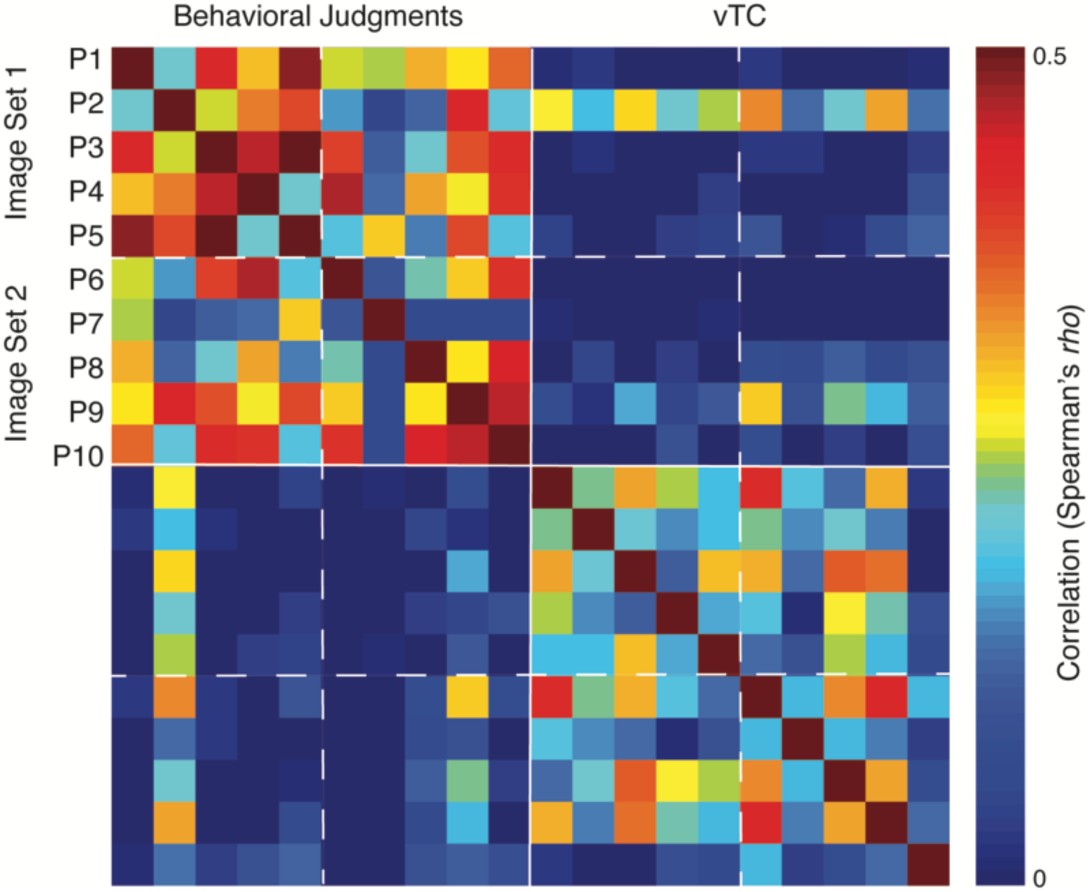
Comparison of individual participant RDMs. At the individual participant level correlations between RDMs for behavioral similarity judgements and fMRI responses in vTC (lower left, upper right quadrant) were weaker than those within each experimental measure (upper left and lower right quadrant). Thus, an individual’s behavioral RDM tended to be more correlated to another subject’s behavioral RDM than to their own vTC RDM.

### Structure of category representations: Hierarchical Clustering

To investigate the nature of the category representational structure, we conducted hierarchical clustering analyses (see Materials and Methods). For behavioral similarity judgements, a group of clear and intuitively meaningful clusters emerged, including clusters that appear to reflect ‘urban landscapes’, ‘transportation’, ‘humans’, ‘household items’, ‘animals/insects’, and ‘natural scenes’ (Figure 5A, left). The first branching point in the dendrogram separates animals/insects and natural scenes from all other categories. Thus, animal categories (e.g. farm animals, wild animals) were not grouped with people (i.e. by animacy), but rather were grouped closest to natural objects and scenes (e.g. fire, flowers, beaches). Human categories (e.g. adults, older adults, kids, sports, and body parts) were grouped most closely to people-related objects (e.g. human food, airplanes, trains, bikes) and people-related places (e.g. living rooms, kitchens). These results suggest that behaviorally, participants tended to group images into manmade (including humans) and natural categories (including animals).

**Figure 5:**
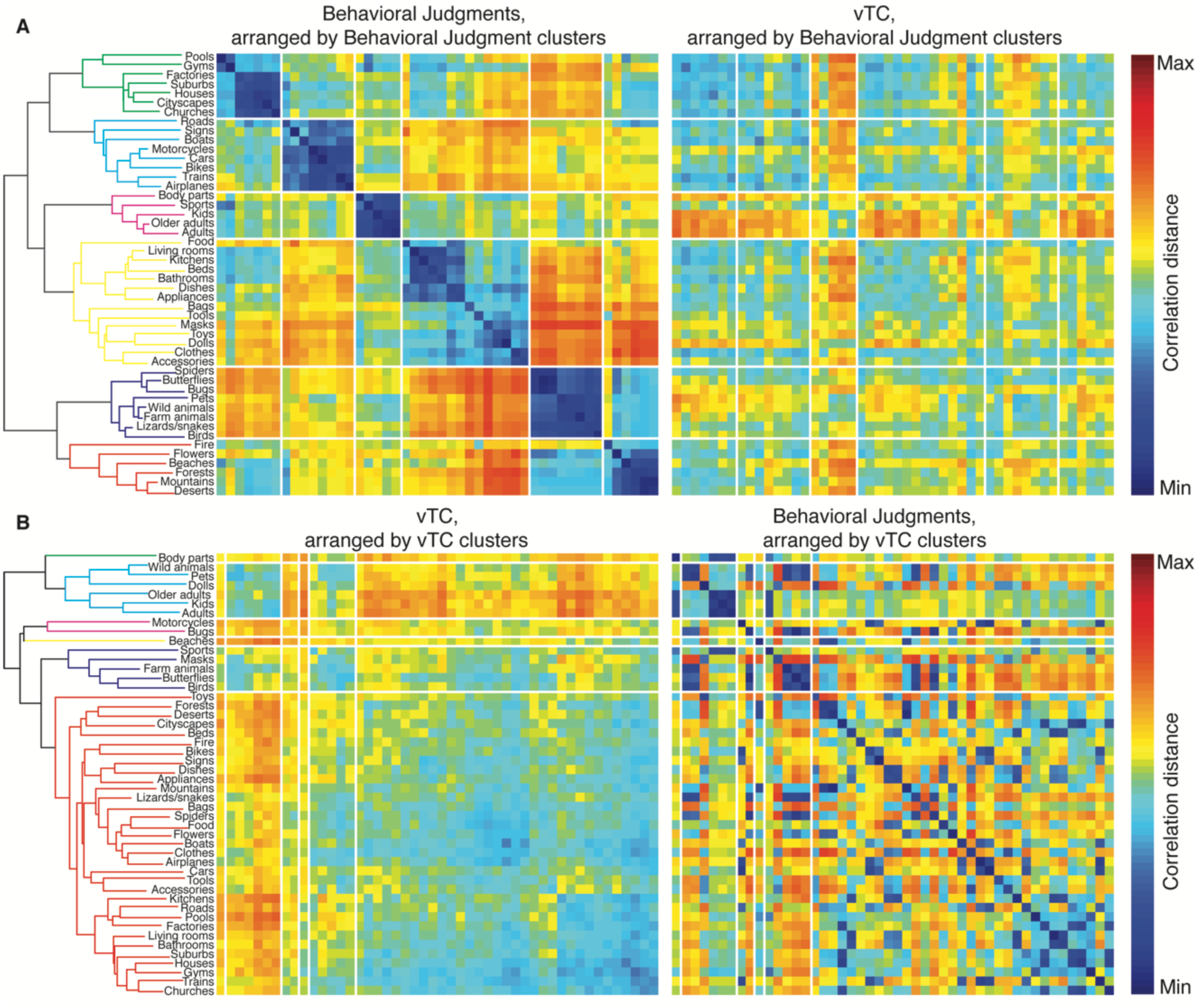
Hierarchical clustering of behavioral and vTC RDMs. A) Hierarchical clustering of behavioral similarity judgments. RDMs for behavior (left) and vTC (right) arranged in the behavioral dendrogram order. B) Hierarchical clustering of vTC dissimilarity. RDMs for vTC (left) and behavioral judgments (right) arranged in the vTC dendrogram order. Dendrogarms are colored according to the top six clusters and the white lines on the RDMs show the boundaries between these clusters.

In contrast, however, hierarchical clustering based on data derived from vTC revealed a relationship between categories that is much harder to characterize (Figure 5b, left). In general, it appears that some categories containing stimuli with faces and/or bodies (e.g. wild animals, pets, dolls, older adults, kids, adults) were represented as similar to one another and distinct from all other categories in vTC, a division that is reflected in the first branching point of the dendrogram. However, there is not a clean grouping of images containing faces and/or bodies from all others since some categories containing faces or bodies (e.g. farm animals, masks) were not contained in the same cluster. In terms of a possible animate/inanimate distinction, it is clear that many animate categories (e.g. lizards/snakes, spiders) were clustered with inanimate categories (e.g. food, flowers, boats, etc.).

Applying the hierarchical clustering orders to the behavioral and vTC RDMs (Figure 5A, B right) highlights the differences between the behavioral and vTC RDMs. When the behavioral clustering order is applied to the vTC RDM, very little structure is present except for the grouping of the categories of kids, adults and older adults, which were relatively more similar to each other than any other categories except for farm animals, wild animals and pets. This suggests some similarities in the representation of kids, adults and older adults between behavior and vTC. When the vTC clustering order is applied to the behavioral RDM, many of the clusters in the behavioral data become fragmented, but some groupings remain. For example the grouping of older adults, kids and adults is clear as well as that of farm animals, butterflies and birds.

In sum, the hierarchical clustering reveals no evidence for a separation of animate and inanimate categories in either the behavioral or the vTC RDM. Moreover, we observe clear differences in the representational structure of the behavioral and vTC RDMs, with more discrete clustering in the behavioral compared to the fMRI domain. The one clear consistency between the behavioral and vTC RDMs is the grouping of the kids, adults and older adults categories. In the next section, we consider whether the differences between the behavioral and vTC RDMs reflect the particular ROI chosen for the fMRI data.

### Beyond the vTC ROI

To investigate whether the weak relationship observed between the behavioral and vTC RDMs reflects the *a priori* choice of ROI, we identified a number of other ROIs in visual cortex and conducted a series of exploratory analyses to determine if any of these regions are more closely correlated with the representational structure that emerged in the behavioral similarity judgments.

First, we defined a series of new ROIs using either independent functional localizers and anatomical constraints (see Methods and Figure 6A). In particular, we examined i) a high level visual region in lateral occipitotemporal cortex (lOTC), analogous to the vTC, incorporating face-, scene-, and object-selective regions, ii) functionally-defined category-selective regions, including both face-selective (FFA and OFA) and scene-selective (PPA and OPA) regions in ventral temporal and lateral occipital cortex, respectively and iii) early visual cortex (EVC) ROIs (combining V1-V3) subdivided into a dorsal (dEVC) and ventral (vEVC) division. We compared the RDMs for each ROI across Image Set 1 and Image Set 2 and also correlated them with the RDM for behavioral judgments.

**Figure 6:**
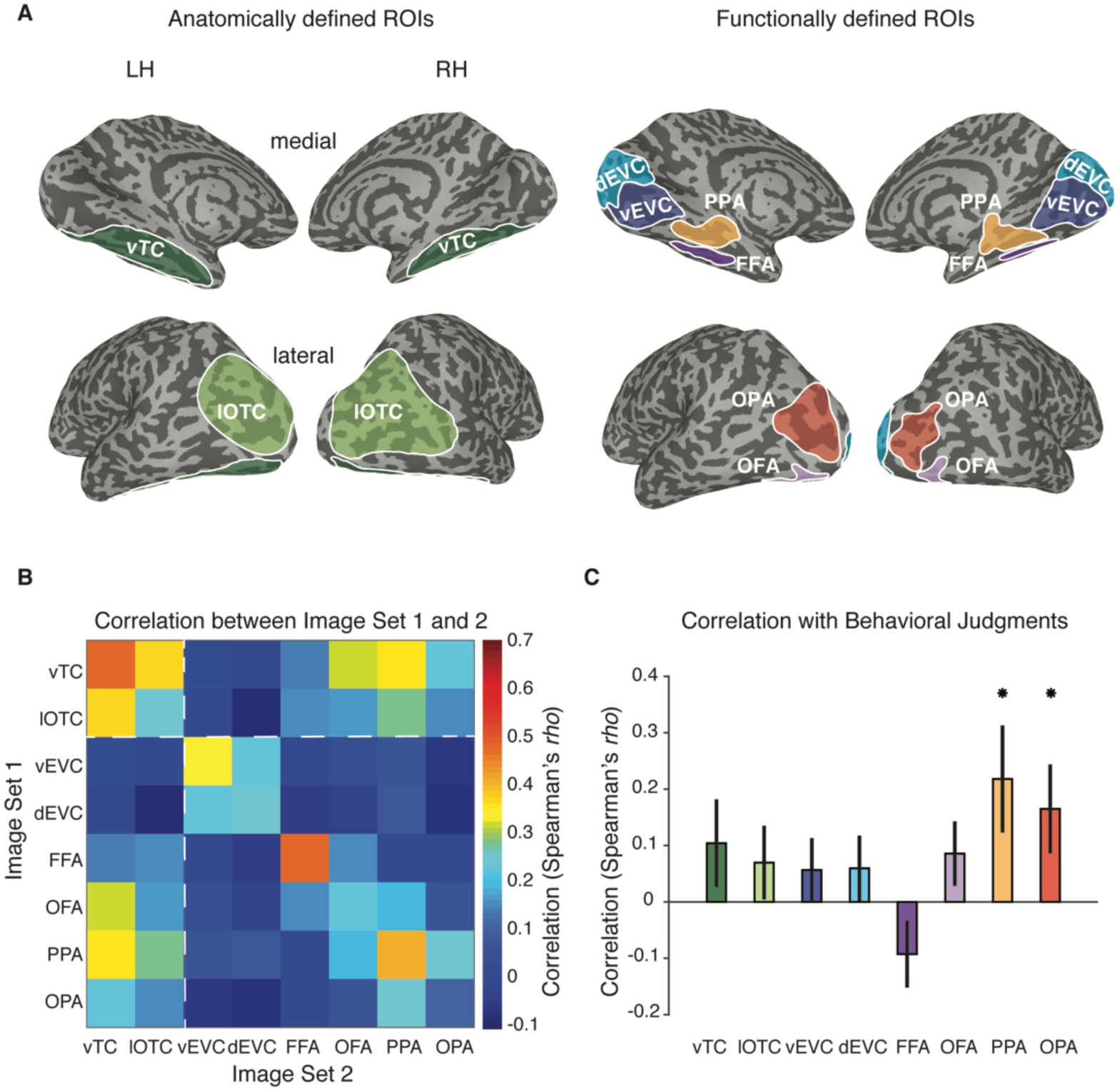
Comparison of multiple visual cortical ROIs. A) Anatomically (left) and functionally defined (right) ROIs. Anatomical and category-selective ROIs were defined in each individual participant. Early visual cortex ROIs were defined at a group-level in an independent set of participants. B) Correlation between the RDMs for each region of interest. Correlations are computed between participants viewing Image Set 1 and those viewing Image Set 2. ROIs included high-level visual cortex on the ventral (vTC) and lateral (lateral occipitotemporal cortex, lOTC) surfaces, dorsal and ventral early visual cortex (dEVC, vEVC), face-selective (OFA, FFA) and scene-selective (OPA, PPA) cortex. Correlations within a ROI were higher on the ventral compared to the lateral/dorsal cortex for all pairs of regions. C) Correlation between the average behavioral RDM and the RDM for each ROI. ^*^ Significant correlations (FDR-corrected) relative to zero (two-tailed) as assessed with a permutation test (n = 10,000). Error bars reflect the standard deviation of the bootstrap distribution of correlation values. The strongest correlation was observed in PPA and the weakest in FFA. Note that the multiple comparisons correction renders the correlation between behavior and vTC reported in our earlier analyses no longer significant.

The diagonal of the ROI comparison matrix (Figure 6B) indicates the reliability of the representational structure across image sets and participants. There are clear differences in the strength of the correlations for the different ROIs. In general, reliability was higher for the ventral compared to the dorsal ROIs (vTC vs. lOTC, vEVC vs. dEVC, FFA vs. OFA, PPA vs. OPA). Further, the representational structure differed across ROIs. For example, the representational structure in the EVC ROIs was very different from that observed in the higher-level ROIs. The vTC ROI, which we used in our analyses so far, varied in its relationship with the other ROIs, showing highest similarity with PPA and lOTC, and lowest with dEVC and vEVC.

For behavior, we compared the RDM for each ROI with the behavioral similarity RDM. PPA showed the strongest correlation (*rho* = 0.22, CI = [0.08, 0.46], *p* < 0.0001) followed by OPA (*rho* = 0.16, CI = [0.07, 0.38], *p* < 0.0001) (Figure 6C), although these correlations were again much weaker than the correlation of the PPA RDM across image sets (*rho* = 0.41, CI = [0.28, 0.59], *p* = 0.0002). The weakest relationship was observed for FFA, which actually showed a trend towards a negative correlation (*rho* = −0.09, CI = [−0.13, 0.10], *p* = 0.06), despite showing a strong positive correlation across image sets (*rho* = 0.49, CI = [0.38, 0.63], *p* < 0.0001).

Second, we conducted an exploratory searchlight analysis to examine any other brain areas that might show a relationship to the representational structure of the stimuli that emerged in behavioral similarity judgments. Our slice prescription included all of occipital, temporal and parietal cortex, but not frontal regions. The strongest brain-behavior correlation emerged in areas corresponding to scene-selective regions PPA and OPA (Figure 7), as well as a medial parietal region that seems to correspond to a third scene-selective region (medial place area, MPA, also referred to as retrosplenial complex, RSC (Epstein, 2008; Silson et al., 2016b).

**Figure 7:**
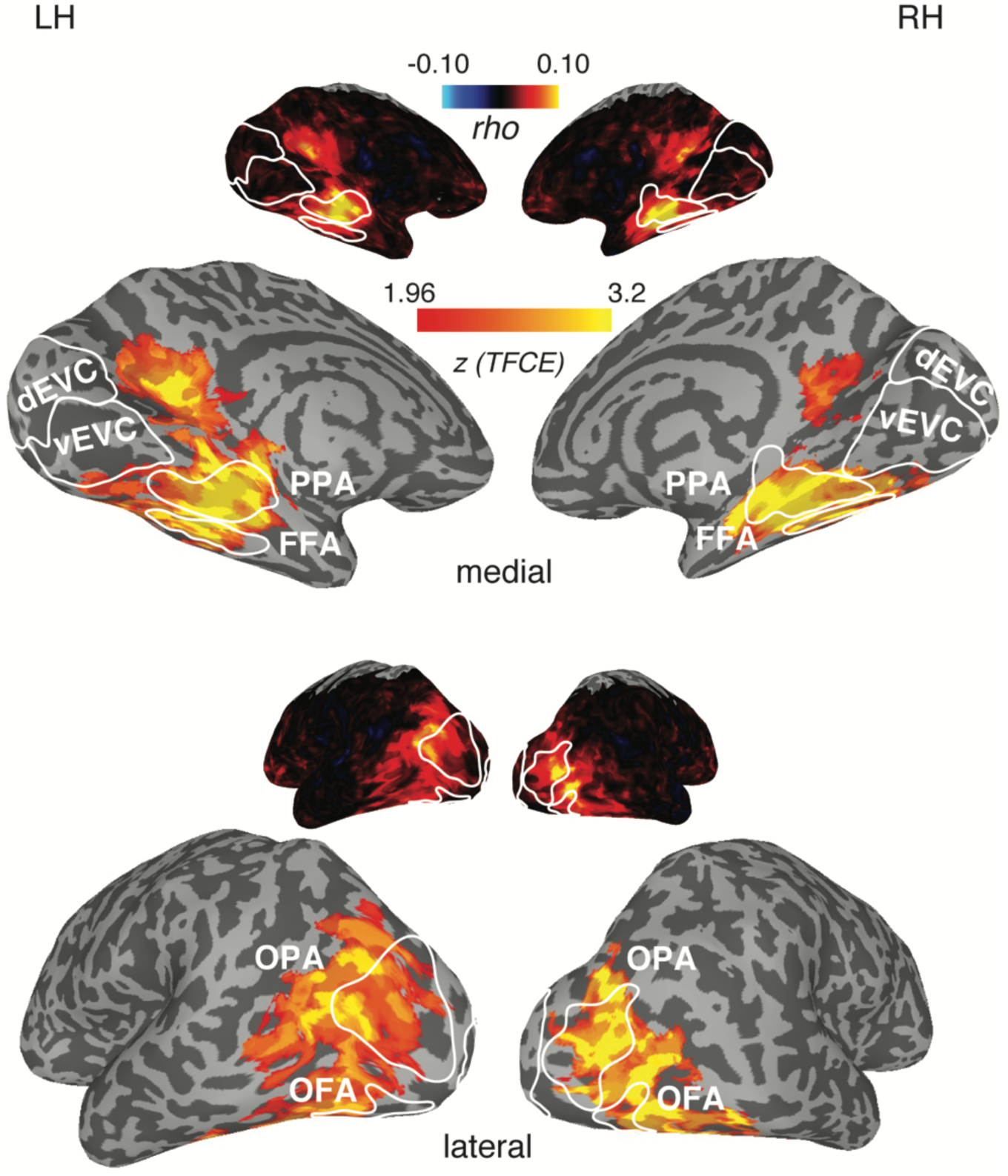
Behavioral RDM searchlight results. The strongest correlations with the behavioral RDM were observed in scene-selective regions OPA and PPA. There was also a strong correlation in medial parietal cortex that likely corresponds to a third scene-selective region, MPA (medial place area). Small brains show the unthresholded correlation values and large brains are cluster-corrected for multiple comparisons using Threshold-Free Cluster Enhancement (thresholded on z = 1.94, corresponding to two-sided *p* < 0.05). Group-level results are overlaid on the freesurfer reconstruction of one example participant, with the corresponding functionally-defined ROIs highlighted in solid white lines.

Taken together, these data indicate the strongest relationship between the representational structure of behavioral similarity judgments and fMRI responses is in scene-selective cortex, particularly PPA, followed by OPA, while the weakest relationship was observed for FFA. This could be considered surprising, given that the one clear consistency between the behavioral judgments and fMRI responses in vTC (a large ROI that encompasses both PPA and FFA) appeared to reflect a grouping of the adults, kids and older adults categories, which are image categories that FFA responds strongly to, but PPA does not. To further explore the origin of this correspondence, we next examined the representational structure in PPA and FFA and their relation with the behavioral dissimilarity in more detail.

### Representation of human categories in PPA and FFA

Hierarchical clustering (Figure 8) indicated that both PPA and FFA contained an early branching of a cluster that included adults, kids and older adults, similar to the larger vTC ROI. However, in PPA, this cluster also included body parts, while in FFA this cluster also included sports (which typically contained people) and dolls. Further, inspection of their respective RDMs (Figure 9A) revealed some clear differences in representational structure. While for both FFA and PPA the categories of adults, kids and older adults showed strong dissimilarity with most other categories (presumably resulting in them being grouped separately in a cluster in both cases), in FFA these categories were also similar to one another, as well as to pets, wild animals and farm animals. In contrast, PPA showed no such grouping by similarity of these categories, instead exhibiting high similarity between urban scenes such as houses, cityscapes and churches, categories that were highly dissimilar from one another in FFA.

**Figure 8:**
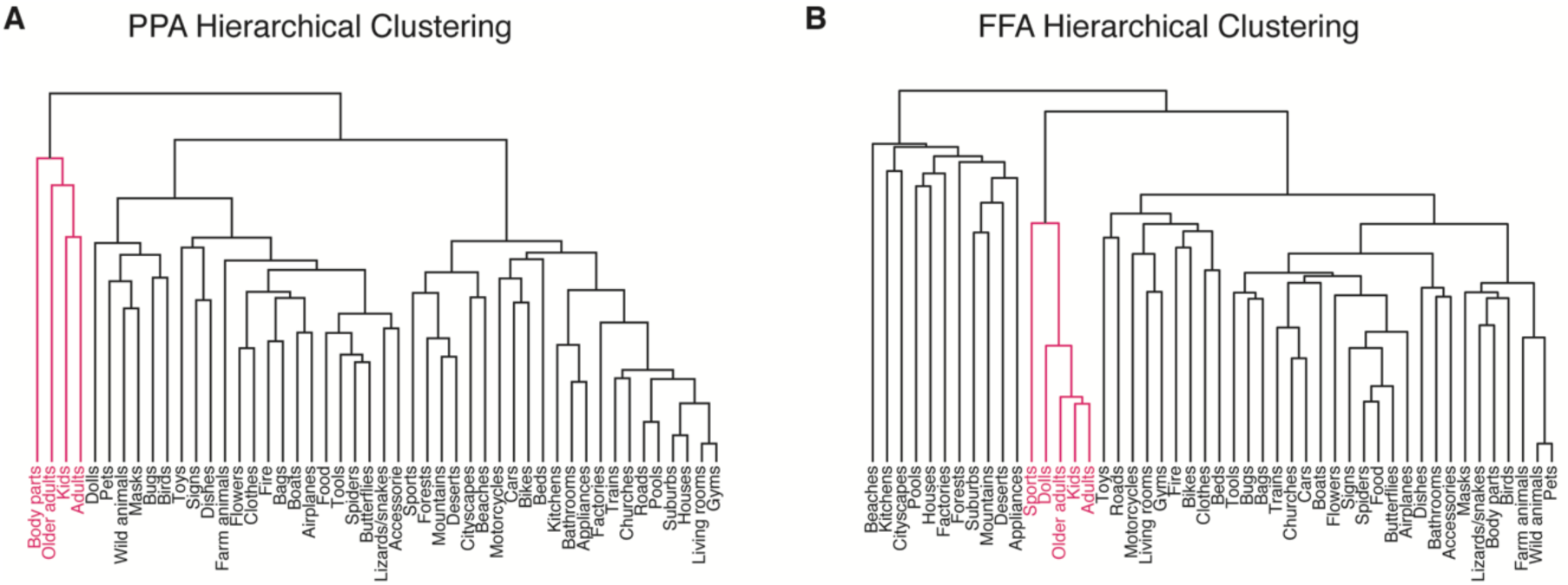
PPA versus FFA: hierarchical clustering. A) Hierarchical clustering of representational dissimilarity in scene-selective PPA indicated the presence of a face- and body-selective cluster (first branch) containing the categories adults, kids and older adults, as well as body parts. B) Hierarchical clustering of face-selective FFA indicated a face-selective cluster (second branch) containing adults, kids and older adults, as well as sports (which typically included people) and dolls.

**Figure 9:**
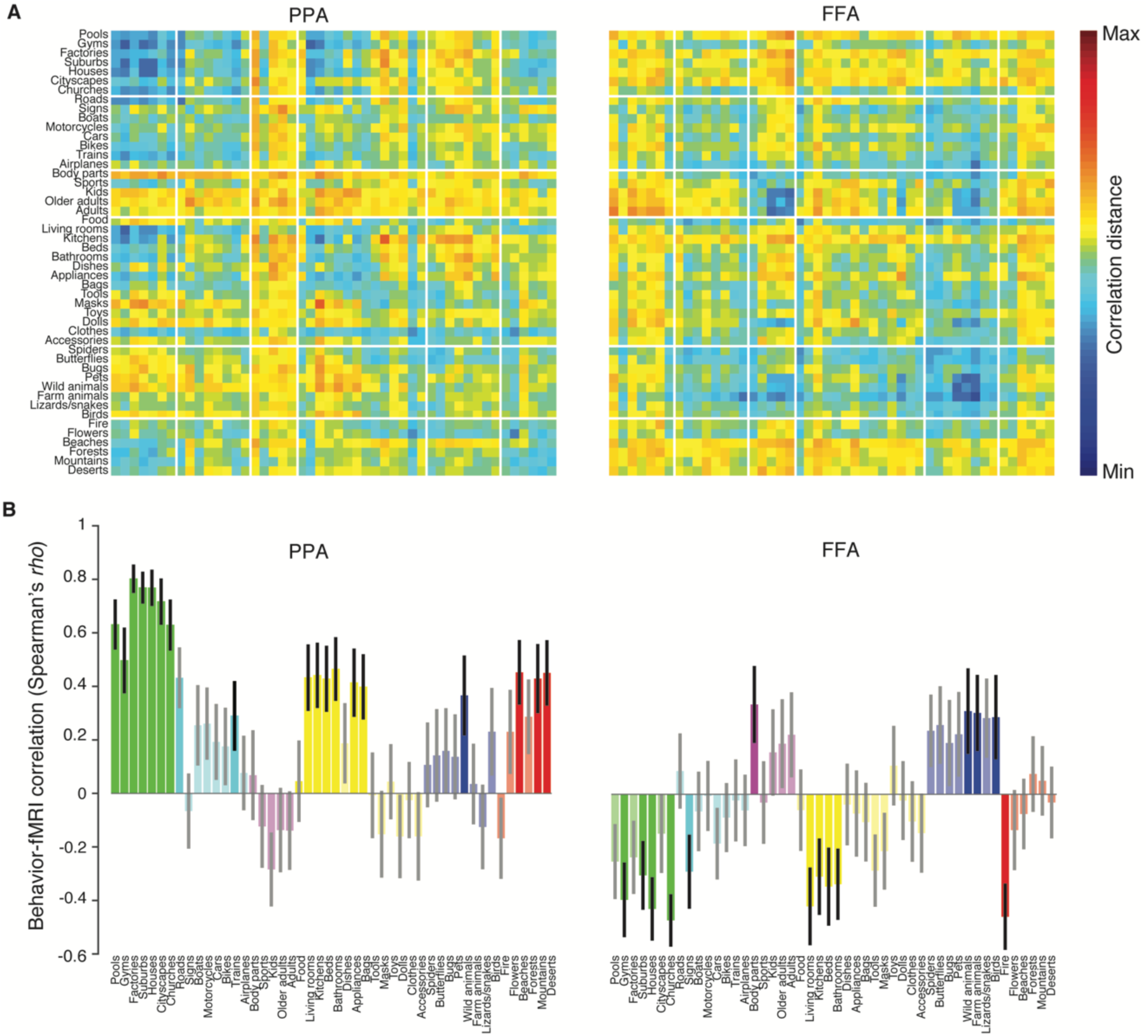
PPA versus FFA: RDMs and individual category correlation with behavior. A) RDMs of PPA and FFA arranged in the behavioral clustering order. Superimposed white lines indicate the clusters derived from the behavioral judgments RDM (see Figure 5A). B) For each category, correlations were computed between PPA (left) or FFA (right) dissimilarity and behavioral dissimilarity (Spearman’s *rho*). Individual correlations are color-coded by the clusters derived from behavioral judgments. Significant correlations are depicted as opaque bars, while non-significant correlations are transparent. Significance was assessed using a permutation test with 10,000 permutations per category (*p* < 0.05, two-tailed). Error bars reflected the standard deviation of the bootstrap distribution of correlations (10,000 bootstraps).

This difference between the PPA and FFA RDMs was further highlighted when the correlation between PPA or FFA and behavioral judgments was computed for each category separately (Figure 9B). High correlations indicate that the category was similarly represented in the fMRI and behavioral RDM, while low or negative correlations indicate differences in the representational structure. For PPA, most categories showed a positive correlation, with the strongest correlations for urban landscapes such as factories, houses and cities. The lowest correlations were observed for categories containing humans or faces such as adults, kids, masks and dolls. In contrast, in FFA, most of the correlations were negative, indicating a striking difference in the representational space for most categories. The strongest positive correlations were observed for categories containing people and for animals. Collectively these analyses suggest that PPA and FFA each capture different aspects of the behavioral similarity judgements.

In sum, comparisons of regions beyond the vTC ROI suggest that representational structure was most reliable for ventral regions, with clear differences in representational structure between regions. Out of all ROIs examined, scene-selective regions correlated best with behavior, and this observation was supported by the searchlight results. However, relative to the reproducibility within the fMRI domain, the magnitude of the fMRI-behavior correlations remained relatively weak. The separation of the kids, adults and older adults categories that we observed for vTC was evident in hierarchical clusters obtained for both PPA and FFA. However, for PPA, the correlation with behavior was driven by non-face categories, while FFA only correlated weakly with behavior for those categories and exhibited limited correspondence for other categories. Collectively, these results suggest that neither ROI fully captured the representational structure reflected in the behavioral judgments. To better understand what is being represented in behavioral judgements and fMRI responses, we next considered a third domain of representation: computational modeling.

### DNN comparisons with fMRI responses and behavioral judgments

In light of previous reports showing a correspondence between DNNs and both behavioral judgments and brain activity measurements in humans and non-human primates, we next examined to what extent DNN representations were able to explain the representational structure observed in our current data. In particular, given the discrepancy between our fMRI and behavioral measurements, we were interested to determine which of the two domains corresponded more strongly with the DNN representations.

We created RDMs based on DNN representations for individual layers of an 8-layer, off-the-shelf pre-trained DNN (see Materials and Methods), separately for Image Set 1 and Image Set 2. Dissimilarities were calculated as the correlation distances between the vectorized responses across all units within a given layer. Similar to the behavioral and fMRI measurements described above, representational structure within each DNN layer (Figure 10A) was reproducible across image sets, increasing gradually from lower to higher layers (Image Set 1 versus Image Set 2, all *rho* = [0.21, 0.62], all *p* < 0.0001). For comparisons with representational structure in the behavioral judgments and fMRI, responses we averaged the RDMs across the two image sets separately for each layer. We then compared the representational structure of each layer with the RDMs for behavioral judgments and a number of fMRI ROIs (Figure 10B).

**Figure 10:**
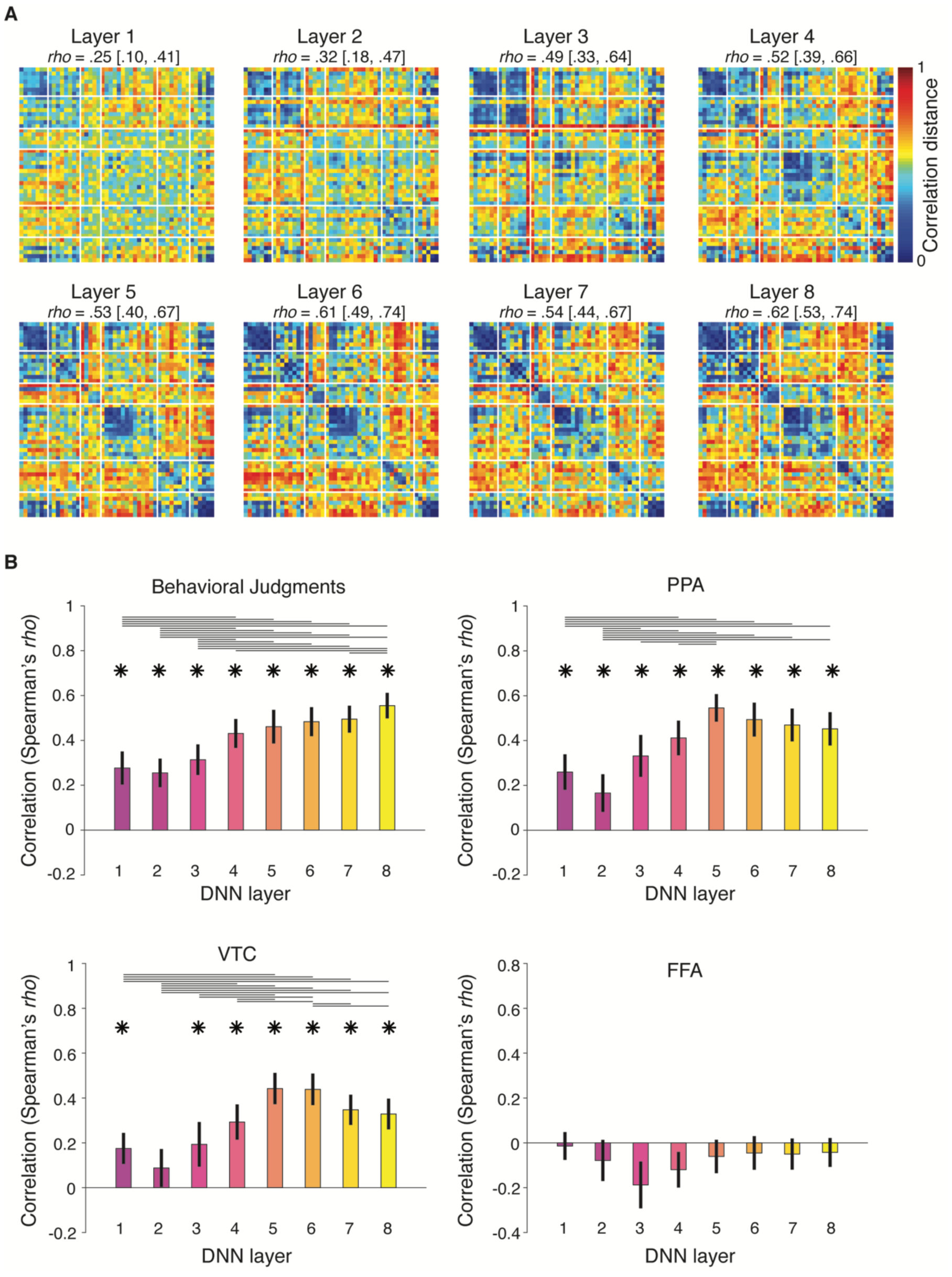
DNN representations correlate with brain and behavior. **A)** RDMs (correlation distances) for each of the 8 layers of the DNN, ordered based on the hierarchical clustering of the behavioral RDM. Superimposed white lines indicate the cluster derived from the behavioral judgments RDM (see Figure 5A). The between set correlation values above each RDM (*rho* [95% CI]) increase with layer number, reflecting increased reproducibility of representational structure for higher DNN layers. **B)** Correlation of each individual layers with behavior, vTC, PPA and FFA. ^*^ significant correlations (FDR-corrected) relative to zero (two-tailed) as assessed with a randomization test (n = 10.000). Horizontal lines indicate significant differences (FDR-corrected) between correlations (two-tailed) as assessed with bootstrapping (n = 10.000). Error bars reflect the standard deviation of the mean correlation, obtained via a bootstrapping procedure (see Methods).

For behavior, we observed a consistent correlation with the DNN that gradually increased with higher layers, culminating in the highest correlation for layer 8 (*rho* = 0.56, CI = [0.46, 0.69], *p* < 0.0001). In contrast, the highest correlation with PPA was found for layer 5 (*rho* = 0.55, CI = [0.44, 0.68], *p* < 0.0001); while its correlation also gradually increased from layer 1 to 5, higher layers did not differ significantly from layer 5. A similar pattern of results was observed for the larger vTC ROI (highest correlation with layer 5: *rho* = 0.44, CI = [0.32, 0.60], *p* < 0.0001). In contrast, none of the DNN layers exhibited a significant correlation with FFA, whose correlations instead appeared to trend negatively (all *rho* = [−0.18, −0.01], all p > 0.05), similar to the relationship between FFA and behavior.

These results demonstrate that higher-level DNN representations are reproducible across image sets and, surprisingly, are correlated with *both* the behavioral and the brain measurements in PPA and vTC, with relatively high maximal correlations for both domains (around *rho* = 0.55). However, behavioral and fMRI representational similarity differed in terms of which layer correlated more strongly. For behavioral judgments, higher layers invariably resulted in increasing correspondences with behavior, all the way to the top-most layer that is closest to the output (layer 8). In contrast, correlations with fMRI measurements in high-level cortex regions increased up to mid-level layer 5, only to plateau or even decrease again for subsequent layers.

This result suggests that additional computations carried out in the fully-connected layers (6-8) are important to explain human behavioral judgments, but not fMRI responses, which map more strongly onto representations contained in the mid-to-high-level convolutional layers.

## Discussion

We compared the representational similarity of behavioral judgments with those derived from fMRI measurements of visual cortex for a set of naturalistic images drawn from a range of object and scene categories. While the representational structure for each type of measurement was reproducible across image sets and participants, there was surprisingly limited agreement between the behavioral and fMRI results. While the behavioral data revealed a broad distinction between manmade (including humans) and natural (including animals) content, with clear sub-groupings of categories sharing conceptual properties (e.g., transportation: roads, signs, airplanes, bikes), the fMRI data largely reflected a division between images containing faces and bodies (e.g. kids, adults, older adults, body parts) and other types of categories, with sub-groupings that were very heterogeneous. This discrepancy was not due to the specific cortical regions chosen, and even the region showing the strongest correlation with behavior (scene-selective PPA) exhibited quite distinct representational structure from that observed for behavioral judgments. An off-the-shelf DNN appeared to explain both the behavioral and fMRI data, yet the behavior and fMRI data showed maximal correspondences with different layers, with fMRI responses mapping more strongly onto middle levels of representation compared to behavior. Collectively, these results demonstrate that there is not a simple mapping between multi-voxel responses in visual cortex and behavioral similarity judgments. Below, we discuss three potential explanations for this divergence.

### 1) Visual versus conceptual information

One possibility is that while the fMRI data reflect the visual properties of the stimuli, behavioral similarity judgments reflect conceptual structure that goes beyond those visual properties. Such a view is consistent with prior studies demonstrating that low-level visual properties contribute to responses in high-level regions of visual cortex (Watson et al., 2017; Groen et al., 2017). Our comparison with the DNN representations seem to support this suggestion, with fMRI most related to layer 5 and behavior corresponding most strongly to layer 8, consistent with prior studies reporting a peak correlation between scene-selective cortex and layer 5 in similar networks (Bonner and Epstein, 2017; Groen et al., 2018; but see Khaligh-Razavi and Kriegeskorte, 2014). The type of DNN layer may be an important factor as layers 1-5 are convolutional and contain ‘features’ that can be visualized (Zeiler and Fergus, 2014) and are still spatially localized in the image. In contrast, layers 6-8 perform a mapping of those features onto the class labels used in training. Thus the later DNN layers contain a potentially more fine-grained categorical representation that better matches behavior of human observers, while the fMRI responses correspond to an earlier stage of processing where visual features relevant for categorization are represented at a coarser level.

Others have suggested, however, that hierarchical visual models (e.g. HMax, DNN) do not capture semantic or conceptual information and that an additional level of representation is required (Clarke and Tyler, 2014; Clarke et al., 2015; Devereux et al., 2018). However, this view tends to discount the covariance between visual features and conceptual properties as well as co-occurrence statistics (e.g. a banana and an orange are much more likely to occur in an image together than a banana and a motorcycle). Indeed, the correspondence we observed between the higher levels of the DNN and behavioral similarity judgments, which appear to reflect fine-grained groupings of conceptually-related stimuli, suggests that a significant amount of conceptual information can be captured by a feedforward visual model.

While we focused on visual cortex, it has been reported that conceptual representations are reflected beyond visual cortex in perirhinal cortex (Devereux et al., 2018; Martin et al., 2018). However, our searchlight analysis demonstrated the strongest correlations between fMRI and behavioral similarity measures in scene-selective regions and did not highlight perirhinal cortex. Our slices included occipital, temporal and parietal cortices but not prefrontal cortex, so it is possible that a stronger correspondence between the fMRI and behavior could emerge there.

### 2) Organization of representations in the cortex

In this study we compared behavioral similarity judgments with representations in regionally-localized brain regions using multi-voxel patterns. In this context, there are two important factors to consider, namely i) the scale and ii) the distribution of information representation in the cortex.

First, multi-voxel patterns may primarily reflect the large-scale topography of cortex rather than more fine-grained representations (Freeman et al., 2011). In high-level visual cortex, there are large-scale differences across the vTC reflecting the categorical distinction between faces and scenes that overlap with an eccentricity gradient (Hasson et al., 2002) and variation according to the real-world size of objects (Konkle and Oliva, 2012). These considerations are consistent with the general grouping we observed in the fMRI data that seemed to reflect a separation of images with faces and bodies from all other images. An alternative approach to using multi-voxel patterns is to model feature-selectivity at the individual voxel level (Naselaris et al., 2011). While this approach might be more sensitive to more fine-grained selectivity, it is striking that studies using this approach have primarily revealed smooth gradients across visual cortex that largely seem to reflect the large-scale category-selective organization (Huth et al., 2012; Wen et al., 2018) with evidence for a limited number of functional sub-domains (Çukur et al., 2013, 2016).

Second, the behavioral similarity judgments revealed apparent conceptual groupings that likely reflect multiple dimensions on which the images could be evaluated. A strong correspondence between a localized cortical region and the behavioral similarity judgments would suggest that all those dimensions are represented in a single region (i.e. a ‘semantic hub’; Patterson et al., 2007). However, we found no such region in our searchlight analysis, suggesting that if it does exist, it likely lies outside of visual cortex. Alternatively, conceptual knowledge may be distributed across multiple regions with each representing specific object properties (Martin, 2016) and there is some fMRI evidence for distributed semantic representations (Huth et al., 2012). However, we also failed to observe a good correspondence with behavior in our vTC ROI, which include a large proportion of high-level visual cortex. While it is possible that some differential weighting of the response across this region may have led to a better fit with the behavioral response, this possibility only further highlights the difficulty in mapping between the response of high-level visual cortex and behavior.

### 3) Task differences

The behavioral task required participants to compare simultaneously presented stimuli and make explicit similarity judgments, but an unrelated fixation cross task was performed during fMRI. It is thus possible that during fMRI participants processed the images differently, resulting in a different representational space (Mur et al., 2013) and a more explicit and involved fMRI task might have yielded more similar representations across tasks. However, while task has been reported to have a strong impact on behavioral representations (Schyns and Oliva, 1999; Harel and Bentin, 2009; Bracci et al., 2017a), fMRI studies have found limited effects of task on representations in vTC (Harel et al., 2014; Bracci et al., 2017a; Groen et al., 2018; Hebart et al., 2018). Instead, task effects appear to be much more prevalent in parietal and frontal regions (Erez and Duncan, 2015; Bracci et al., 2017a; Vaziri-Pashkam and Xu, 2017). In fact, the relative inflexibility of representations in vTC compared to behavior further highlights the difficulty in directly mapping between them.

#### Representation of animacy

One striking aspect of our results is that contrary to previous work (Kriegeskorte et al., 2008; Naselaris et al., 2012; Mur et al., 2013; Sha et al., 2015) we did not observe a clear separation of animate vs. inanimate categories in either behavioral or fMRI representational similarities. Instead, in behavior, images were initially grouped according to a broad division between man-made (including humans) and natural categories (including animals). With fMRI, we observed a separation of face and body categories from all others. This difference with the prior literature could reflect a broader sampling of categories in our study or the use of backgrounds rather than segmented objects presented in isolation (Kriegeskorte et al., 2008; Sha et al., 2015). However, evidence for an animate distinction has been reported even with a large sampling of natural scenes (Naselaris et al., 2012). Alternatively, it is also possible that what has been termed animacy in previous studies primarily reflects the presence of face or body features and not animacy *per se*. Indeed, a recent study found that animate objects (e.g. cow) and inanimate objects that looked like an aimate object (e.g. cow-shaped mug) are represented similary in vTC (Bracci et al., 2017b).

## Conclusion

By comparing behavioral similarity judgments with fMRI responses in visual cortex across a range of object and scene categories, we find that while there is a correlation between fMRI and behavior, particularly in scene-selective areas, the structure of representations is strikingly different. Further, while both the behavior and the fMRI data correlate well with DNN features, the modalities best matched different levels of representation. Collectively, these results suggest that there is not a simple mapping between localized fMRI responses and behavioral similarity judgments with each domain capturing different visual properties of the images.

## Acknowledgements

We thank Susan Wardle and Martin Hebart for helpful discussion and comments on earlier versions of this manuscript, Ed Silson for help with the ROI definitions, and Steven Scholte for help implementing the DNN analyses. This research was supported by the Intramural Research Program of the US National Institute of Mental Health (ZIAMH 002909), Clinical Study Protocol 93-M-0170, NCT00001360. The authors declare no competing financial interests.

**Author Contributions**
MLK, DJK and CIB designed the study. MLK and IIAG performed the research. MLK, IIAG, AS and DJK analyzed the data. MLK, DJK, IIAG, AS and CIB wrote the paper.

